# Influence of deterministic and stochastic processes on microbial community assembly during aerobic granulation

**DOI:** 10.1101/356303

**Authors:** Raquel Liébana, Oskar Modin, Frank Persson, Enikö Szabó, Malte Hermansson, Britt-Marie Wilén

## Abstract

Aerobic granular sludge is an energy efficient and compact biofilm process for wastewater treatment which has received much attention during the last decades and is now being implemented in full-scale. However, the factors involved in microbial community assembly during formation of granules are poorly understood and little is known about the reproducibility in treatment performance and community structure. Here we show that both deterministic and stochastic factors exert a dynamic influence during microbial community assembly into granular sludge. During granulation, the microbial communities in three replicate sequencing batch reactors followed similar successional trajectories of the most abundant taxa and showed similar dynamics in diversity. Deterministic factors dominated the assembly of the most abundant community members as the microbial community transitioned from floccular to granular form. Stochastic factors mostly affected rare members of the communities and caused the microbial community structure to diverge in one of the reactors; however, this did not have an impact on the treatment performance. This demonstrates that the reactor function and the dynamics of the most abundant community members are in fact reproducible during the formation of aerobic granules.

## Introduction

The structure of microbial communities depends on complex and dynamic ecological mechanisms. Studying the compositional differences between two different communities (β-diversity) is useful in order to understand the underlying ecological mechanisms that have shaped the communities, since it provides a link between biodiversity at the local scale (α-diversity) and at the regional scale (γ-diversity) [1, 2]. Vellend (2010) developed a conceptual framework, modified by Nemergut *et al* (2013), where the ecological mechanisms responsible for community assembly are framed into four fundamental processes: selection, dispersal, diversification and drift. Selection results from deterministic factors that modify the community structure according to the environmental conditions, differences in fitness between individuals and microbial interactions. Dispersal results both from deterministic and stochastic factors which enable the movement and establishment of organisms among communities. Diversification results mainly from stochastic factors which generate genetic variation. Drift refers to stochastic changes due to birth, death and reproduction. These four processes act simultaneously in natural ecosystems and their influences vary in time and space [4, 5].

Generally, the design and development of wastewater treatment systems is hindered by the lack of understanding of the ecology of the microbial communities on which they rely [6, 7]. The importance ascribed to the different ecological mechanisms for shaping the microbial community composition in wastewater treatment bioreactors vary in the literature. Some studies report similar microbial communities in replicate reactors, such as membrane bioreactors [8], anaerobic digesters [9–11], microbial fuel cells [12] and sequencing batch reactors [13, 14], due to selection caused by the reactor environmental conditions. On the contrary, diverging trajectories have been reported in replicate microbial electrolysis cell reactors [15], sequencing batch reactors [16] and anaerobic digesters [17, 18], due to the roles of stochastic factors. However, deterministic and stochastic factors are not mutually exclusive but exert a combined effect on the microbial community assembly and need to be assessed in conjunction [4, 5].

Wastewater treatment bioreactors are engineered to select for the functional groups needed for water purification. Also, the reactor operational conditions are adjusted to cultivate the microorganisms in the desired aggregate modes. The microbial communities in wastewater treatment are either assembled as flocs in activated sludge or as biofilms on different substrata. Free-floating spherical biofilms, so called granules, combine many of the properties of these two growth modes [19, 20], which has gained increasing interest and implementation in recent years [21]. The possibility to remove carbon, nitrogen and phosphorous simultaneously is one of the most important attributes of granular sludge, which renders highly compact and less energy demanding wastewater treatment [22]. The granules are compact and spherical suspended biofilms with a diameter of approximately 1–3 mm obtained in defined reactor conditions of high up-flow air velocity, large shear forces, large temporal variation of electron donors and -acceptors causing feast-famine operation and short settling time to select for well settling biomass [23, 24]. Despite well-established methods for granule cultivation, detailed studies on the microbial community ecology in aerobic granules are scarce and the ecological processes underpinning the microbial community assembly during granulation are poorly understood [25].

Laboratory experiments in wastewater treatment are often performed in one reactor due to practical reasons. Hence, conclusions regarding reactor performance and microbial community structure are generally drawn from single reactors operated at different conditions [26–29]. It is unclear how reliable these conclusions are, especially regarding complex processes such as microbial community assembly. Furthermore, laboratory scale wastewater studies that enable controlled environmental conditions and tests of reproducibility are valuable test-benches for studies of microbial ecology [30]. To better understand he microbial community assembly and reproducibility of granule formation, we have used amplicon sequencing to investigate the succession during the transition from floccular to granular sludge in three replicate reactors seeded with the same inoculum and operated at identical conditions for 35 days. At the same time, the function of the microbial communities was monitored. We hypothesized that similar communities would develop in all three reactors if deterministic factors were dominating, whereas the opposite would support the importance of stochastic factors. We also investigated possible shifts in the importance of deterministic/stochastic factors for community assembly during the operation period.

## Material and methods

### Experimental set-up

Three identical sequencing batch reactors (6 cm diameter and 132 cm height) with a working volume of 3 L were operated for 35 days at room temperature (20–22°C) in 4-hour cycles (0:05 h filling, 0:55 h anaerobic, 2:53 h aerobic, 0:02 h settling, 0:05 h withdrawal) with a volumetric exchange ratio of 43%. The settling time was gradually decreased to promote granulation and at the same time allow retention of slow growing organisms (Figure S1), and the aerobic phase was adjusted accordingly to permit even 4-hour cycle lengths. Air was provided from the bottom of the reactors with porous diffusers (pore size 1 μm) at a flow rate of 2.5 L min^−1^ and superficial up-flow velocity of 1.5 cm s^−1^. The reactors were inoculated with aerobic/anoxic activated sludge from a full-scale wastewater treatment plant (Gryaab AB, Gothenburg, Sweden). Synthetic wastewater was pumped in at the bottom of the reactor with a chemical oxygen demand (COD) to nitrogen ratio of 100:25 (3 kg COD m^−3^d^−1^ and 0.75 kg NH_4_-N m^−3^d^−1^) and pH of approximately 7 and consisted of 1.6 g l^−1^ NH_4_CH_3_COO, 12.5 mg l^−1^ MgSO_4_-7H_2_O, 15.0 mg l^−1^ CaCl_2_, 10.0 mg l^−1^ FeSO_4_-7H_2_O, 22.5 mg l^−1^ K_2_HPO_4_, and 1 ml l^−1^ micronutrient solution (0.05 g l^−1^ H_3_BO_3_, 0.05 g l^−1^ZnCl_2_, 0.03 g l^−1^ CuCl_2_, 0.05 g l^−1^ MnSO_4_.H_2_O, 0.05 g l^−1^ (NH_4_)6Mo_7_O_24_.4H_2_O, 0.05 g l^−1^ AlCl_3_, 0.05 g l^−1^ CoCl_2_.6H_2_O, and 0.05 g l^−1^ NiCl_2_). Biomass samples (100 ml) were withdrawn from the reactors daily for analysis. The replicate reactors were operated identically.

### DNA extraction, amplification and sequencing

Samples for microbial community analysis were collected from the reactor three times per week. Total genomic DNA was extracted using the FastDNA Spin Kit for Soil (MP Biomedicals) following the manufacturer’s protocol. The V4 region of the 16S rRNA gene was amplified in duplicates, using the forward primer 515’F (5’-GTGBCAGCMGCCGCGGTAA-3’) and the reverse primer 806R (5’-GGACTACHVGGGTWTCTAAT-3’), indexed according to Kozich *et al* (2013). The PCR reactions were conducted in a 20 μl reaction volume using 17 μl of AccuPrime Pfx SuperMix (Life Technologies), 1 μl of genomic DNA (20 ng template), and 1 μl each of the forward and reverse primers (10 μM). The PCR reactions were carried out in a Biometra T3000 Thermocycler with the following thermal cycling parameters: initial denaturation at 95°C for 5 min, followed by 30 cycles of denaturation (95°C, 20 s), annealing (50°C, 15 s) and elongation (68°C, 60 s), and finished by a 10 min final elongation at 68°C. PCR products were purified with the MagJET NGS Cleanup and Size Selection Kit (Thermo Scientific) and the DNA concentration was measured using a Qubit 3.0 fluorometer (Invitrogen), using the dsDNA HS assay kit (Invitrogen). The PCR products were pooled in equimolar amounts, the concentration and size was confirmed by TapeStation 2200 (Agilent Technologies) and sequencing was performed with a MiSeq using reagent kit v3 (Illumina). To increase the confidence in the obtained results, a second MiSeq sequencing run was performed on the same pool. Raw sequencing data were deposited in the NCBI Sequence Read Archive (SRA), BioProjectID PRJNA472243, accession number SRP148672.

### Sequence processing and data analysis

Raw sequence reads were processed following the UNOISE pipeline [32, 33] with USEARCH v.10 [34]. Briefly, paired-end reads were merged with a maximum number of mismatches of 15 (5% of the read length) and a minimum of 80% of identity in the alignment. Merged reads were filtered (maximum expected error per read of 0.5), dereplicated (default settings) and denoised (minimum abundance of 4), resulting in zero-radius operational taxonomic units (OTUs). OTUs were taxonomically classified with the SINTAX algorithm [35] based on the MiDAS database v.2.1 [36] and analysed in R version 3.4.1 (http://www.r-project.org). The dataset was rarefied, subsampling each sample to 17395 sequences. Basic R functions were used to perform Wilcoxon signed-rank tests and to calculate Pearson correlation coefficients. Non-metric multi-dimensional scaling (NMDS) ordination and heatmaps were created using the R package ampvis [37]. Constrained analysis of proximities (CAP) was performed using the R package vegan [38]. The sequences were aligned with the R package DECIPHER [39] and a maximum likelihood tree was generated with FastTree 2 software [40] using the GTR+CAT (General Time Reversible with per-site rate CATegories) model of approximation for site rate variation and computation of Gamma20-based likelihood.

α-diversity for each sample was calculated using numbers equivalents (Hill numbers) [41]. We refer to these indices as naïve diversity (^q^ND) because the phylogenetic distance between OTUs are not considered. The parameter q is the diversity order. At a q of 0, all OTUs are considered equally important and, hence, ^0^ND is the sample richness (i.e. the number of OTUs in a sample). For higher values of q, more weight is put on abundant OTUs. We also calculated diversity numbers which take the sequence dissimilarity into account using the framework described by Chiu and Chao (2014). The metric, here referred to as ^q^PD (phylogenetic diversity), represents the total effective phylogenetic distance between all OTUs in a sample. For all pairwise comparisons of samples, β-diversity was obtained using the same calculation framework. The β-diversities were converted into dissimilarity indices (^q^βdis) constrained between 0 (two identical samples) and 1 (two samples with no shared OTUs) [42, 43]. The parameters ^q^βdis_ND_ and ^q^pdis_PD_ are based on the naïve and the phylogenetic β-diversity values, respectively.

### Null model analysis

Null models were used to analyse the taxonomic- and phylogenetic successional turnover between consecutive samples in the reactors. Taxonomic turnover was estimated with Raup-Crick-based measures based on Bray-Curtis dissimilarities (RC_bray_), calculated with the R package vegan as in Stegen *et al* (2013). Shortly, a null distribution (999 randomizations) of expected Bray-Curtis dissimilarities was created for each pair of communities, which was compared to the empirically observed Bray-Curtis dissimilarities. The RC_bray_ index is calculated by adding the number of times the null distribution is greater than the empirical Bray-Curtis distribution to half the number of times the null and the empirical distribution are equal. Then this count is normalized to a value between −1 and 1. A negative value means that the two communities share more taxa than expected by chance whereas a positive value means that they share less taxa. Values > |0.95| were considered statistically significant. Values < |0.95| indicates that the taxonomic turnover between the community pair was not different from the null expectation and therefore, influenced by stochastic factors [5].

The R package PICANTE [45] was used to calculate the β nearest-taxon index (βNTI), which measures if the phylogenetic turnover is greater or less than the null expectation. For this, the β mean nearest-taxon distance (βMNTD), which measures the mean phylogenetic distance between the most closely related OTUs in two communities, was first calculated based on relative abundance data, as in Stegen *et al* (2013). In each iteration (999 randomizations) the OTUs were moved randomly across the tips of the regional pool phylogeny and the resulting phylogenetic relationships were used to calculate the βMNTD_null_. The βNTI was calculated as the difference in standard deviation units between the observed βMNTD and the mean of βMNTD_null_ in the pairwise sample comparisons. A negative βNTI value means that the samples are more phylogenetically close to each than expected by chance and a positive value means that the two samples are more phylogenetically distant from each other. Previous studies have found that short phylogenetic distances between OTUs, quantified using βMNTD and βNTI, reflect ecological distances [44, 46, 47]. Pairwise comparisons with βNTI > |2| were considered statistically significant. Values < |2| were not significantly different from the null expectation, which indicate that stochastic factors influenced the phylogenetic turnover [5].

### Analytical methods

Concentrations in the effluent of total organic carbon (TOC) and total nitrogen (TN) were measured with TOC-TN analyser 87 (TOC-V, Shimadzu, Japan). Total suspended solids and volatile suspended solids in the in the reactor (MLSS, MLVSS) and in the effluent (effTSS, effVSS) were measured according to standard methods. Microscopy was performed using an Olympus BX60 light microscope (Olympus Sverige AB, Solna, Sweden) with particle size assessment by measuring the diameter of 10 random flocs/granules using the CellSens (Olympus) software.

## Results

### Granulation of the sludge and reactor performance

The sludge started to granulate around day 7, when small granules emerged (Figure S2) and the mean particle sizes of the reactor biomass increased similarly in the three reactors (Figure S3). At day 25, the granulated biomass dominated in the reactors and at the end of the experiment (day 35), the granules had an average diameter of 1.5 mm. The three reactors also followed similar patterns over time in suspended solids concentrations in the reactor and in the effluent (Figure 1A-B), although R1 had significantly lower reactor concentrations (MLSS and MLVSS) than R2 and R3 (p<0.05) (Table S1). Removal of TOC and TN was similar among the reactors (Figure 1, C-D). TOC removal was above 90% since day 4, with an average of 95% for R1 and R3 and 96% for R2. The average TN removal was above 50% for the three reactors.

**Figure 1.**
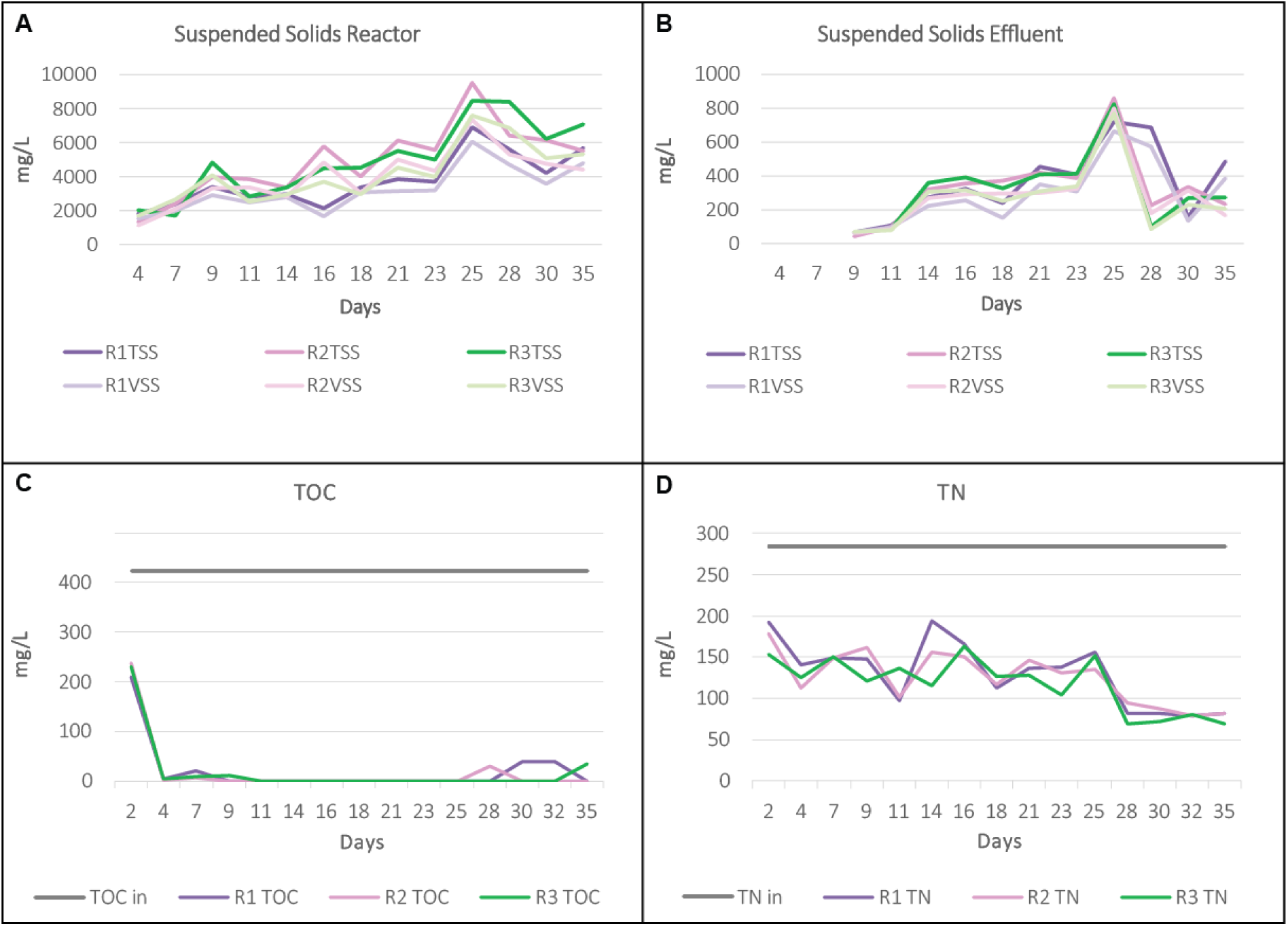
Sludge properties and reactor performances over time for R1 (purple), R2 (pink) and R3 (green). A, reactor suspended solids concentrations; B, effluent suspended solids concentrations; C, TOC concentrations in the influent and effluent; D, TN concentrations in the influent and effluent.

### Temporal variation of abundant taxa

The microbial community composition was highly dynamic during the transition from floccular to granular sludge. The seed sludge, mainly composed of *Betaproteobacteria* (38% of total reads), *Alphaproteobacteria* (14%), *Sphingobacteria* (11%) and *Clostridia* (7%) at class level, changed rapidly after inoculation to a community dominated by *Gammaproteobacteria* (70%) with *Acinetobacter* as the dominant genus during the first 7 days (Figure 2, S4). Thereafter, *Betaproteobacteria* dominated in the three reactors, with high abundances of the genera *Thauera, Acidovorax, Simplicispira, Comamonas, Curvibacter* and *Alicycliphilus* (Figure 2). From days 21–25, *Actinobacteria* dominated until the end of the experiment, with *Corynebacterium* sp. as only representative of this class, having more than 30% of the total reads in the three reactors. Many taxa that were rare or undetected in the seed sludge became abundant during the granulation, e.g. *Corynebacterium* sp., *Brevundimonas* sp., *Pedobacter* sp., *Fluviicola* sp. and *Bacteriovorax* sp., while other taxa that were common in the seed sludge decreased drastically in relative abundance, e.g. *Rhodoferax* sp., *Rhodobacter* sp., *Pseudorhodobacter* sp. and *Hydrogenophaga* sp.

**Figure 2.**
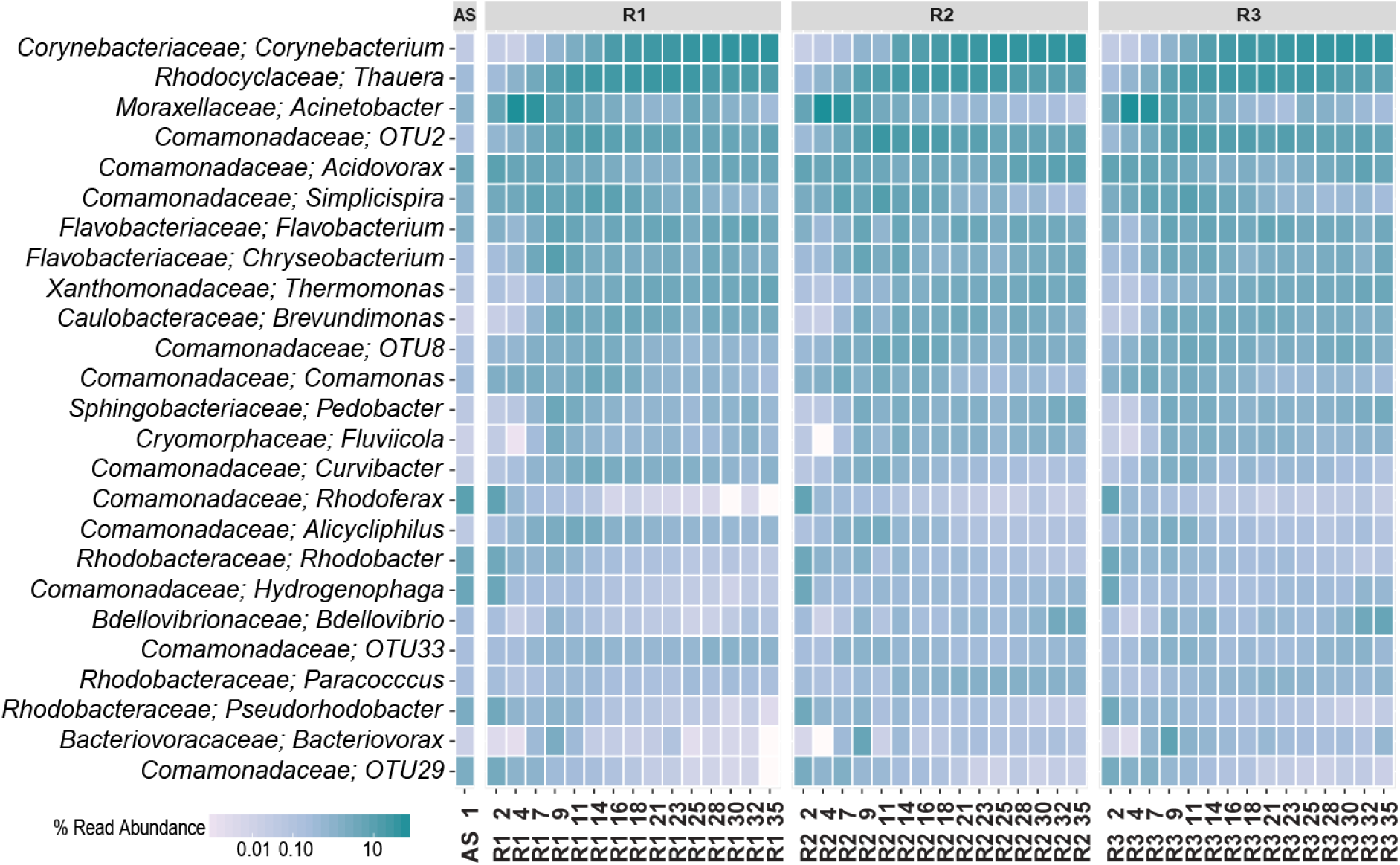
Temporal variation of the 25 most abundant genera as percentages of reads in the replicate reactors (R1, R2 and R3) and the seed sludge (AS), from days 1 to 35.

### Microbial diversity and dynamics during granulation

The three reactors followed the same trends in α-diversity (Figure 3A-C). After inoculation, there was a drop in diversity, followed by a peak around day 9 and then a continuous decrease. R1 differed somewhat from R2 and R3 during the dynamic period after the inoculation and during days 21–35 when ^0^ND was consistently lower in R1, while ^1^ND and ^2^ND were similar in the three reactors, showing that the rare OTUs were fewer in R1. The observed trends for naïve diversity were very similar to the trends for the phylogenetic diversity (Figure S5A-C).

**Figure 3.**
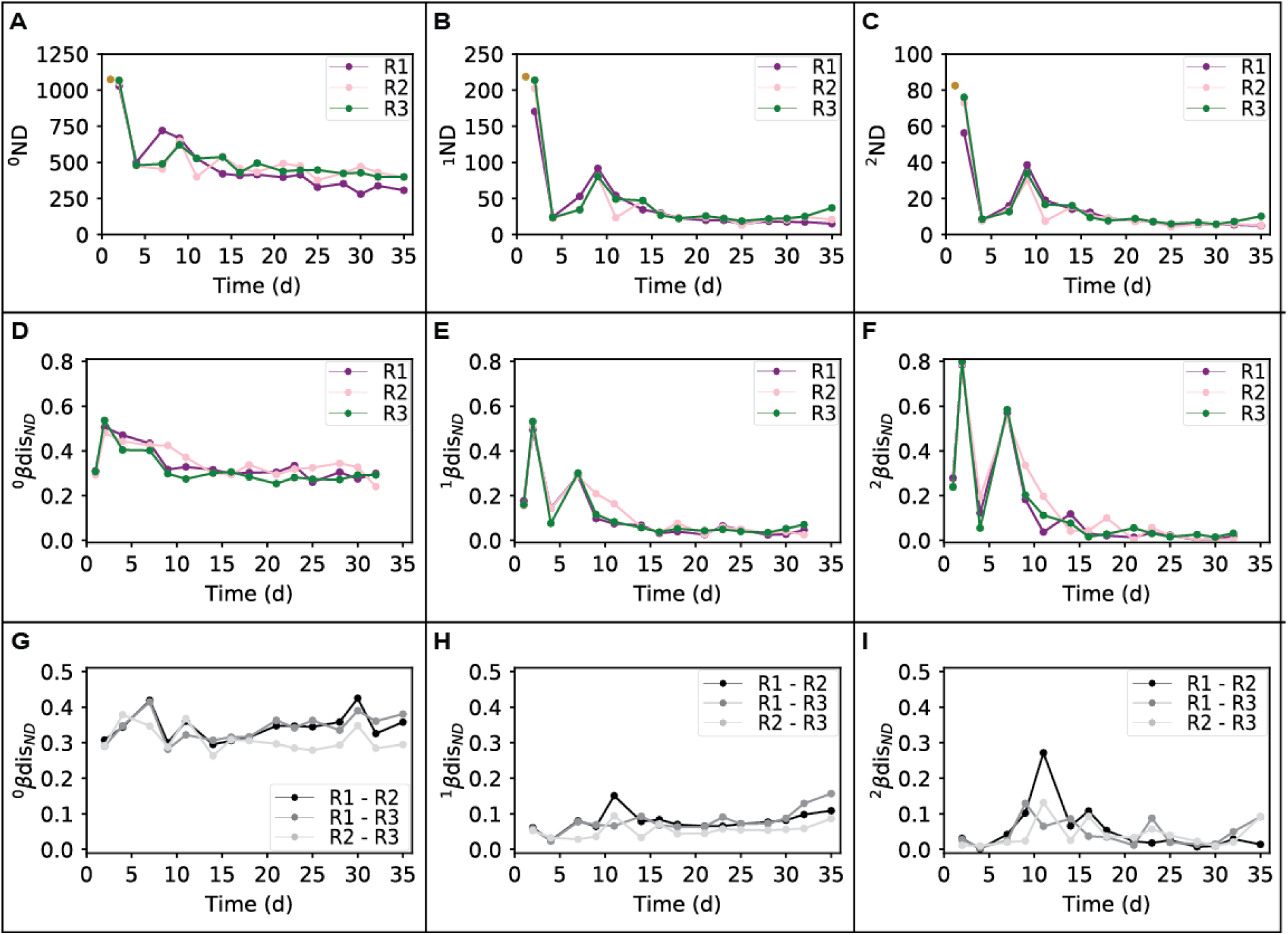
Dynamics of the naïve diversity (taxonomic distance between OTUs) for the replicate reactors. A, B and C: naïve α-diversity; C, D and F: naïve β-diversity over time between two successive sample points; G, H and I: naïve β-diversity over time between the replicate reactors.

Succession, measured as β-diversity between successive sample points, showed that after inoculation there were large temporal dynamics in each reactor with dissimilarities of around 0.5 for ^0^βdisND and even higher peak dissimilarities (0.8) for ^2^βdisND, which gives large weight to abundant OTUs (Figure 3D-F). Thereafter, the dissimilarities between successive samples decreased considerably to values of about 0.3 for ^0^βdisND and 0.05 for ^1^βdisND and ^2^βdisND.

For β-diversity between the reactors (Figure 3G-I), ^0^βdisND, was around 0.3–0.4, showing that 60–70% of the OTUs were shared among the reactors at any given time. ^1^βdisND was around 0.1 and ^2^βdisND around 0.05, i.e. the reactors were more similar to each other when considering the more abundant OTUs. R2-R3 were more similar than R1-R2 or R1-R3 for diversity order 0 and 1, but not for diversity order 2. Also for β-diversity between successive sampling points or between reactors, the naïve diversity patterns were highly similar to the ones for phylogenetic diversity (Figure S5).

NMDS shows initial, rapid changes after inoculation and that R1 and R2 deviated from the other reactors at day 7 and day 11 (Figure 4). From day 14, R2 and R3 converged and evolved similarly after that, whereas R1 diverged. CAP analysis based on Bray-Curtis dissimilarities confirmed R1 to be different to R2 and R3 (Figure S6). Pearson correlation based on CAP showed that 144 out of the 2540 OTUs were significantly different (p<0.05 in the correlation to CAP2 axis, see Table S2) in relative abundance between reactors. 58 OTUs were more abundant in R1 than R2-R3 and 82 OTUs more abundant in R2-R3 than R1. These OTUs, mainly of low relative abundance added up to 8% of the total reads.

**Figure 4.**
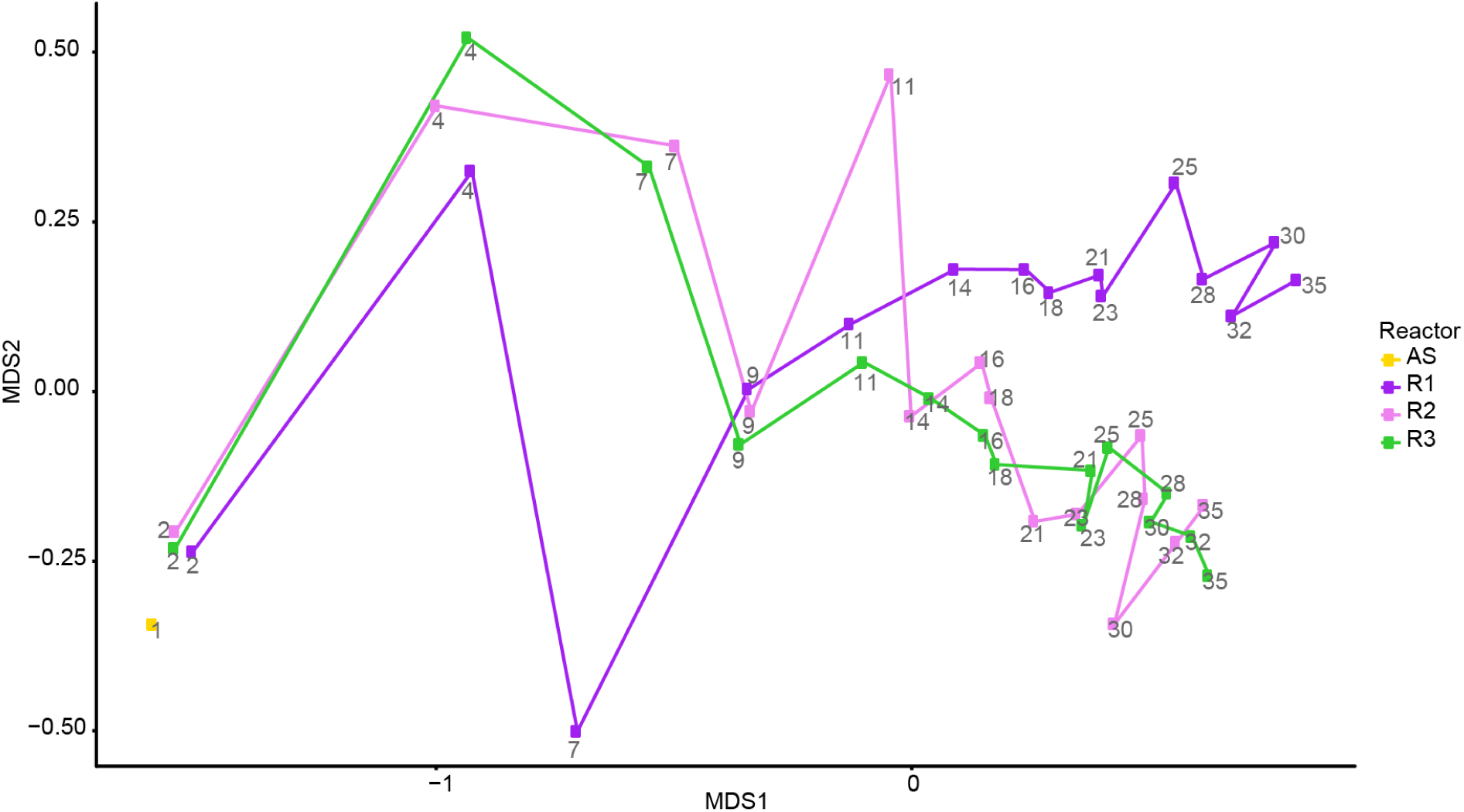
NMDS ordination based on Bray-Curtis dissimilarities after square root transformation. Each line represents the community succession of each reactor, R1 (purple), R2 (pink), R3 (green), and the seed sludge (AS, day 1, yellow). Numbers refer to days after startup. Stress =0.047.

### Null model analysis

We wished to determine the factors behind the shifts in β-diversity over time in each reactor. However, α-, β- and γ-diversity are interconnected so that e.g. a difference in richness between samples may in itself cause a difference in β-diversity between the samples [15, 48]. Null model analysis was therefore used to disentangle turnover due to succession from changes in α- and γ-diversity. We used two different types of null models for this, RC_bray_, based on taxonomic turnover, and βNTI, based on phylogenetic turnover. Turnover between sequential samples from the same reactor would show successional phases. Thus, RC_bray_ and βNTI values lower than the null expectation, would indicate "steady-state” microbial community composition. In R1, RC_bray_ values were close to 1 during the first 9 days (Figure 5A), indicating higher taxonomic turnover than the null expectation, and thereafter turnover remained lower than the null expectation (RC_bray_ < −0.95). R2 an R3 showed a somewhat different trend with turnover closer to the null expectation (RC_bray_ between 0.95 and −0.95) during the first 11 days and after this period, the taxonomic turnover between successive sample points remained lower than expected by chance (RC_bray_ < −0.95). Similar patterns were observed using βNTI, with higher phylogenetic turnover for R1 than for R2 and R3 until day 14, followed by lower turnover than expected by chance in all reactors (Figure S7). RC_bray_ was also used for the fraction of OTUs having significantly different relative abundance between the reactors, as determined by correlations to the CAP axes (Figure S6 and Table S2). For this fraction of the community, the taxonomic turnover between succesive sample points in R2 and R3 gradually increased from day 9 to values not significantly different to the null expectation whereas in R1, the taxonomic turnover remained lower than the null expectation throughout the reactor operation (Figure 5B).

**Figure 5.**
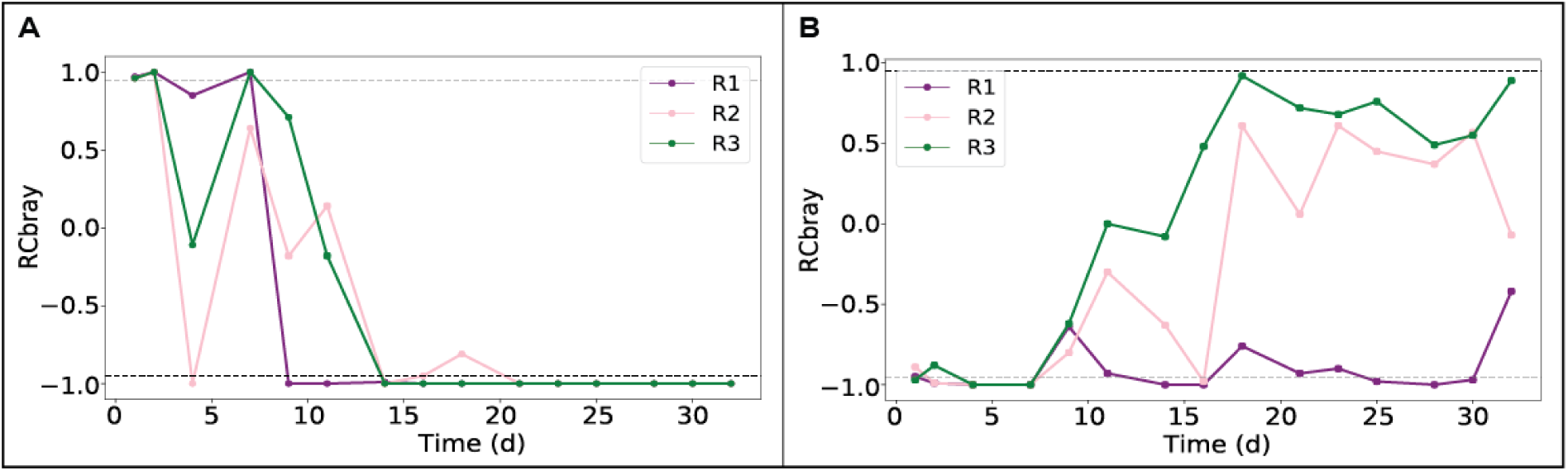
Taxonomic turnover (RC_bray_) in community composition between two succesive sample points of the whole community (A) and of OTUs with a significant p-value (p<0.05) in the Pearson correlation for the CAP2 axis (B) of the replicate reactors.

### Sequencing reproducibility

To ensure that the observed differences between the reactors, mainly caused by OTUs of low abundance, were not caused by limited precision of sequencing, the samples were sequenced twice. Highly similar results were obtained from the two sequencing runs with regard to relative read abundance of major OTUs, NMDS ordination and CAP (Figures S8-S10).

## Discussion

In this study, we investigated how communities of microorganisms assembled over time as granules formed in three replicate reactors for wastewater treatment. Granule growth, successional patterns of taxa, microbial diversity and reproducibility are discussed together with an assessment of the mechanisms that influence the community assembly.

### Succession of taxa during the granulation process

During the early transformation from flocs to granule-like particles, *Acinetobacter* sp. became increasingly abundant in all reactors (Figure 2). *Acinetobacter* sp. has been reported as primary colonizers and bridging bacteria in aggregate/biofilm development [49, 50] and have previously been observed during initial stages of granulation [51]. *Acinetobacter* sp. is a hydrophobic bacterium and produces EPS, which are decisive properties for early stage aggregate formation [52] and likely explains their frequent occurrence during the initial phase.

As the granules started to increase in size, after one week of operation (Figure S3), the microbial communities shifted and bacteria within *Comamonadaceae, Rhodocyclaceae, Flavobacteriaceae, Xanthomonadaceae* and *Caulobacteraceae* became more abundant. Many of these bacteria are associated with granulation [26, 51]. *Thauera* sp., which was particularly abundant in this period (Figure 2), is an important EPS producer commonly found in acetate-fed aerobic granules and is also a well-known denitrifier, as are many of the other common microorganisms in this period [26, 53]. For instance, bacteria within the *Comamonadaceae* family are considered important for acetate-fed denitrification in aerobic granules [54]. Despite this, the total nitrogen removal was incomplete, around 50% until day 28. Thereafter nitrogen removal increased to 70% until the end of the study period, day 35. In this last period, *Corynebacterium* sp. was highly abundant in the reactors (Figure 2). *Corynebacteria* can remove nitrogen both by assimilatory reduction and denitrification [55].

None of the common members of the microbial community in the granules were highly abundant, or even detected in the seed sludge. This is commonly observed in reactor studies [9, 26, 27] and experimental evidence suggests that during succession, rare taxa can be promoted to the dominant fraction of the microbial community following a perturbation or a change of environment [56–58].

### Microbial community assembly during granulation

The microbial communities in the replicate reactors followed similar succession patterns but with a clear divergence of R1 from R2 and R3 (Figure 4). Since the inoculum, feed composition, and reactor operation were the same among the reactors, this suggests that stochastic factors had a role in the community assembly. Let us first consider the two extreme cases of either stochastic or deterministic factors acting alone. With only stochastic factors, i.e. drift, neutral theory predicts that the microbial communities in the three reactors would eventually diverge and the dissimilarity index between reactors would increase and reach unity. On the other hand, if deterministic factors, i.e. selection, were acting alone, the reactors would follow the same trajectory and the dissimilarity index between reactors would be zero. The ecological processes responsible for microbial community assembly can be classified as selection, dispersal, drift, and diversification [3]. Our experimental design excluded dispersal as a major factor because synthetic feed was used. Diversification can most likely also be neglected because of the short time frame of the study (35 days). This leaves selection as the major deterministic process and drift as the major stochastic process affecting community assembly.

During the early start-up, the microbial community shifted rapidly from the inoculum in a similar manner in all reactors (Figure 4) and the phylogenetic successional turnover was relatively high, especially in R1 (Figure S7). The settling time for the biomass was long (Figure S1), thus the microorganisms were not washed out due to their settling properties, but according to their ability to grow at these reactor conditions [59]. Therefore, the changes in the microbial community structure, resulting in a harsh decrease in diversity (Figure 3), were to a large extent caused by the switch from municipal wastewater to a synthetic feed with acetate as carbon source. A switch from a complex substrate to a simple and easily biodegradable substrate would favour specific taxa, and consequently, the biodiversity, both taxonomic and phylogenetic, would decrease [26, 51]. Indeed, the drop in diversity was particularly evident for highly abundant OTUs (^2^ND and ^2^PD, Figure 3, S5) and the dissimilarity between the consecutive days 2 and 4 was also higher for the more abundant OTUs (^2^βdisND and ^2^βdisPD, Figure 3). In addition, the lack of immigration of new taxa (dispersal) to the reactors due the synthetic feed probably contributed to the decrease in α-diversity [60].

The settling time was thereafter drastically reduced, from 30 min to 5 min between days 5 and 11, to promote granulation (Figure S1) and the first granules were observed at day 7. This led to increased α-diversity in the reactors as new, previously undetected, OTUs grew in relative abundance (Figure 3), presumably due to the formation of new niches in the emerging granules. The communities changed rapidly over time, in particular around day 7 (Figure 3), and it was clear that the community assembly took different paths in the three reactors in this phase (Figure 4). The null-models indicated that stochastic factors, here mainly drift, were important for the succession during this dynamic phase (Figure 5, S7). This could be due to the random distribution of microorganisms between aggregates of different sizes of which only the larger and denser ones were retained by sedimentation [59].

From day 11, the decrease in settling time was less dramatic (Figure S1) and the granules increased gradually in size (Figure S3). In this phase, the microbial communities changed more slowly with time (Figure 3, S5) and turnover was lower than predicted by chance by the null-models (Figure 5, S7), suggesting that deterministic forces, i.e. selection, were important. The microbial succession in R2 and R3 followed the same trajectory until the end of the experiment, but R1 diverged (Figure 4). The divergence of R1 was not due to the most abundant OTUs, the community was dominated by the same genera in the three replicate reactors (Figure 2). However, α-diversity (^0^ND) was lower in R1 than R2 and R3 and β-diversity between reactors (Figure 3) showed that the divergence of R1 was mainly caused by rare and intermediate OTUs. A subset of the OTUs were significantly different among the reactors, as assessed by correlation to the CAP2 axis (Figure S6), and these OTUs were of low or intermediate relative abundance (Table S2). For these OTUs, successional turnover was lower than by chance in R1, while in R2 and R3, turnover was higher and the null-model predicted stochasticity (drift) to have larger influence (Figure 5B) Within the same microbial community, selection and drift can affect different subpopulations differently [5]. The less abundant OTUs were likely more susceptible to drift, since many events could result in extinction [4]. Interestingly, predatory bacteria belonging to the genus *Bdellovibrio, Bacteriovorax* and *Peredibacter* were detected in the three reactors at as high as 10.45% maximum relative abundance. *Bdellovibrio* and like organisms (BALOs) prey on gram negative bacteria, with some BALOs being generalist and some specialists in their prey preferences [61]. BALOs have been found in granules [28, 62, 63] and have recently been shown to affect the granular microbial structure [64]. R2 and R3 harboured considerably more BALOs than R1, especially in the later stages of the study (Figure 2 and Figure S8). Also, out of a total of 60 OTUs belonging to BALOs, 11 were significantly lower in abundance in R1 compared to R2 and R3 according to CAP (Table S2). It could hence be hypothesized that BALOs caused a higher turnover in R2 and R3, contributing to the divergence of these reactors from R1.

We tentatively interpret the difference between the reactors in terms of underlying mechanisms: During the early phase the change of feed selected for certain taxa, which caused the biodiversity to decrease (Figure 3). The successional turnover was high, but the microbial communities in replicate reactors were similar (Figure 2 and 4), therefore the acclimation to a new feed seems to be mostly deterministic. A second transitional phase occurred when the settling time was decreased and granules began to emerge. This likely created new niches, which allowed low-abundant taxa, previously below the detection threshold, to increase in abundance resulting in temporarily increased α-diversity (Figure 3). During this phase, stochasticity seemed to cause the microbial community in reactor 1 to take a different path (Figure 4). However, selection eventually led to the same dominating taxa in the parallel reactors (Figure 2). The successional turnover was low during this last period, presumably because selection had led to the development of a microbial community adapted to the specific reactor conditions, but drift caused reactors 2 and 3 to slowly diverge from reactor 1.

### Ecological processes and wastewater treatment

Biological processes for wastewater treatment are often tested in laboratory- or pilot-scale bioreactors. Bioreactors are designed as if they were solely affected by deterministic processes and the treatment process relies on this assumption. But the literature is inconsistent regarding the reproducibility of wastewater biological reactors. Vanwonterghem *et al* (2014) observed that the microbial community structure and dynamics in three replicate anaerobic digesters were highly reproducible, influenced by deterministic processes. Ayarza *et al* (2010) reported the microbial community dynamics to follow similar patterns in four replicate sequencing batch reactors treating synthetic wastewater due to deterministic processes, although, drift caused the reactors to display a differences. Zhou *et al* (2013) demonstrated divergent microbial communities in 14 replicate microbial electrolysis cell reactors treating wastewater due to a dominant role of stochastic processes. Despite highly dynamic microbial communities, stable performance of parallel bioreactors treating wastewater are often found for such general functions as aerobic/anoxic oxidation of organic matter in activated sludge [16, 65–67]. On the other hand, for complex processes that require multispecies cooperation, such as anaerobic digestion or specialized processes such as microbial electrolysis, stochastic variations in community composition has resulted in altered microbial functions [15, 67]. In the present study the differences in the microbial community dynamics were due to differences in the abundance of a small fraction of the total OTUs, which did not have major impact on the reactor functions since the replicate reactors behaved similarly (Figure 1). The same dynamics in dominant OTUs and a higher variability of the less abundant fraction of the community has also been observed in full-scale activated sludge reactors [68]. From the engineering perspective our results are important since they not only confirm the possibility to select for certain dominating taxa by manipulating the reactors conditions, even for a process as common as acetate oxidation, but also provide information about fundamental ecological processes involved in sludge granulation. Moreover, the results presented here suggest that it is possible to draw conclusions about the reactor performance and microbial community dynamics (at least for abundant community members) in experiments with single aerobic granular sludge reactors.

In summary, our study shows that three parallel granular sludge reactors operated identically and inoculated with the same seed sludge behaved similarly, both in terms of treatment performance and microbial community assembly. Deterministic and stochastic processes exerted a dynamic influence during the different successional stages. Deterministic factors shaped the community composition of abundant members whereas stochastic factors mostly affected rare members of the communities. Drift caused the microbial community in one of the reactors to diverge from the other two. Predation seems to have an influence in the divergence of the reactors. Despite these differences in the microbial community, the reactors functions were stable and similar between reactors. More studies dealing with fundamental ecological processes involved in granulation are needed. For instance, it is important to know how perturbations affect the microbial community and the resilience of the system. This knowledge will help to better understand the technology which ultimately will improve its application and management.

## Acknowledgements

This work was supported by FORMAS (Contract no. 245-2013-627), the Swedish Research Council for Environment, Agriculture Sciences and Spatial Planning. The authors acknowledge the Genomics core facility at the University of Gothenburg for support and use of their equipment.

